# CULTURE AND ISOLATION OF BACTERIA ASSOCIATED WITH MEDITERRANEAN CORALS

**DOI:** 10.64898/2026.05.20.726489

**Authors:** Rocío Mozo, Andrea Illa-Oviedo, Javier del Campo

## Abstract

Corals harbor a diverse bacterial community that facilitates adaptation and sustains their health. In coral holobiont research, culture-independent approaches have transformed the existing paradigm. Molecular techniques, such as metabarcoding, revealed a high diversity of previously unrecognized bacterial symbionts. Coral microbiota characterization has relied on these techniques over the last decade, but relying solely on them does not provide a detailed understanding of the dynamics of the coral holobiont complex. Returning to classic microbiological methods and in vitro experimentation can yield novel insights into symbiont roles, physiology, and interactions within the holobiont. Under this premise, we aimed to isolate and culture bacteria from four Mediterranean corals. The recovery of 84 pure bacterial isolates and their initial classification based on the 16S rRNA gene revealed substantial diversity among symbionts amenable to culture. Several isolates represent novel species within relevant genera, such as Vibrio, underscoring the value of culture-based studies. All cultures were cryopreserved to guarantee long-term accessibility for future projects. This represents a key step towards describing the roles of bacteria within the coral holobiont, as cultures enable in-depth morphological and physiological characterization of the symbionts and experimental ecology studies.

## 1. INTRODUCTION

Corals live in association with a complex microbial network conforming the coral holobiont. The coral holobiont is among the best-studied and harbors one of the most diverse microbiotas known to date, comprising bacteria, archaea, viruses, fungi, and protists (del Campo et al., 2020; Rohwer et al., 2002; Rosenberg et al., 2007; Wegley et al., 2007). These microbial partnerships can be determinant for host survival and adaptation to the environment, and together they constitute an ecological and evolutionary unit (Thompson et al., 2015). Bacterial associations were among the first microbial partnerships to be discovered, with early studies developed 30-40 years ago identifying nitrogen-fixing bacteria within the host (Shashar et al., 1994; Williams et al., 1987). This foundational finding led to the development of theories on their role in nutrient acquisition and on the specificity of the bacterial community across different coral species. Nowadays, we know that corals harbor a highly diverse, dynamic, specific, and distinct bacterial consortiums from the surrounding water column (Grottoli et al., 2018; Littman et al., 2009; Rohwer et al., 2002; Speare et al., 2020).

The symbiotic equilibrium between the coral host and its microbial partners is vital for the coral’s health, aiding nutrient acquisition, resilience to environmental stressors, and local adaptation (Osman & Weinnig, 2022; Rosenberg et al., 2007; Webster & Reusch, 2017). The holobiont is an adaptable unit, capable of maintaining homeostasis and generating rapid short-term adaptive responses through shifts in its symbiotic relationships (Reshef et al., 2006; Rosenberg et al., 2007; Webster & Reusch, 2017; Ziegler et al., 2017). However, facing the current climate crisis, extreme and abrupt changes in environmental conditions are disrupting coral holobiont equilibrium, leading to dysbiosis states (such as coral bleaching or coral disease) that drive mass mortality events (Baker et al., 2008; Burke et al., 2023; Hughes et al., 2018). Given this situation, exploring microbial dynamics and describing symbiont roles can help us understand the impact of climate change and guide our conservation efforts.

Although coral-bacteria symbioses have attracted scientific attention for many years, current knowledge is biased due to the intrinsic limitations of the methods used to approach their study. Most of our knowledge derives from barcoding or metagenomic analyses that target specific gene markers, such as 16S rRNA (Grottoli et al., 2018; Keller-Costa et al., 2021). These studies have shed light on the diversity, location within the host, and spatiotemporal variations of these bacterial communities. However, while molecular tools have been helpful in obtaining a broad picture of bacterial associations with corals, relying solely on them is insufficient to develop a comprehensive understanding of the holobiont (Sweet et al., 2021). Many physiological traits and symbiont interrelations can’t be reliably inferred from genetic data alone, and a considerable fraction of microbial diversity remains uncultured and poorly understood.

Genetic data on coral holobionts have become widely available in recent years, but translating these data into functional and ecological insights remains a significant challenge (Robbins et al., 2019; Sweet et al., 2021). To obtain the complete picture, it is essential to complement the existing sequence-based data with in vitro cultures of bacterial symbionts. Access to bacterial isolates will enable their detailed physiological and morphological characterization, which is key to developing a comprehensive understanding of coral holobiont functioning (Li et al., 2022). Such integrative approaches will strengthen our knowledge of host-microbe interactions and their contribution to coral physiology and survivability under changing environmental conditions.

While metabarcoding surveys have extensively characterized the taxonomic composition of bacteria associated with tropical and Mediterranean corals, the culture-based recovery of these microbes remains disproportionately limited relative to sequence-based characterizations. In this context, the present study addresses this imbalance by establishing a culture collection of coral-associated bacteria from four Mediterranean coral species (*Paramuricea clavata, Cladocora caespitosa, Eunicella singularis*, and *Leptogorgia sarmentosa*). Pure symbiont cultures were preserved in a cryobank, providing a valuable resource for future genomic, transcriptomic, and ecophysiological studies.

## 2. MATERIALS AND METHODS

### 2.1 Sample collection and processing

4 coral species (*P. clavata, C. caespitosa, E. singularis*, and *L. sarmentosa*) were sampled via scuba diving surveys along the Spanish east coast (Figure 1), spanning multiple coral colonies to maximize symbiont diversity. Stored in Falcon ® tubes containing seawater from the collection point, samples were either transported to the laboratory for processing or stored at 4 °C. The processing consisted of rinsing the samples with distilled water (to discard not intimately attached bacteria) and cutting them into 3 equivalent fragments. Each fragment was homogenized using a sterile mortar and pestle, and the obtained coral slurry was used as an inoculum for cultivation.

**Figure 1.**
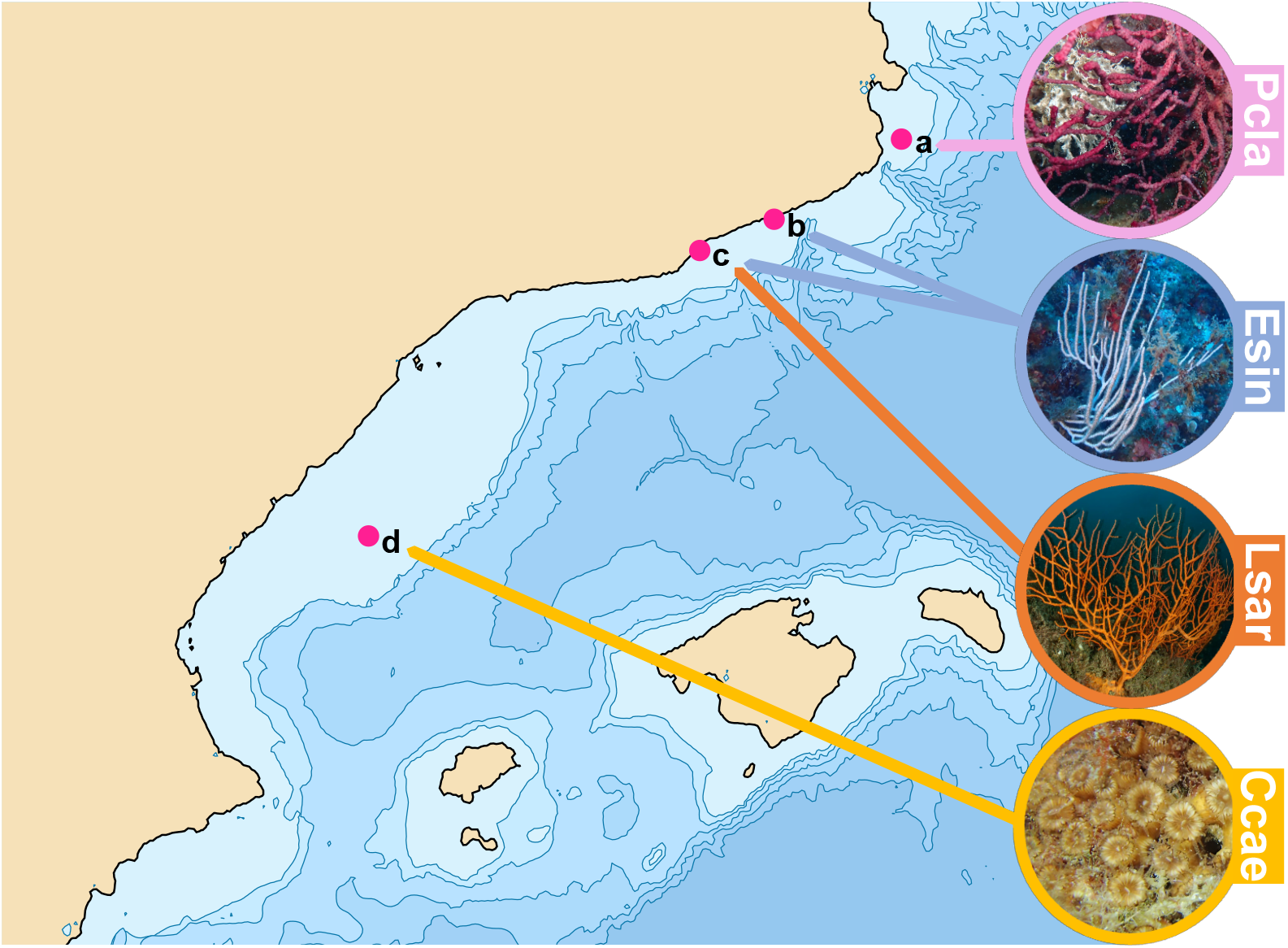
Sampling locations of Mediterranean coral species. Map of the northeastern coast of Spain showing the four sampling sites and the coral species collected at each location: (a) Medes Islands (one *P. clavata* sample), (b) Mataró (three *E. singularis* samples), (c) Sant Sebastià Beach (two *E. singularis* samples and one *L. sarmentosa* sample), and (d) Columbretes Islands (one *C. caespitosa* sample).

### 2.2 Coral-associated bacteria culture and isolation protocol

A series of solid and liquid cultures were performed to achieve pure bacterial cultures. Initially, the coral slurry was streaked onto three different solid media to ensure the culture of the maximum diversity of symbionts: TCBS agar (Difco™), one-half strength Marine Agar 2216 (Difco™) (MA/2, diluting MA in distilled seawater), and MA/2 with coral extract (resuspending coral slurry in MA/2 solution) (Schultz et al., 2022). In order to recruit slow-growing species, morphological characterization of the colonies and isolation were performed repeatedly during three weeks of incubation (in the dark, aerobic conditions, 20-22ºC temperature). The identification of bacterial strains was achieved through morphological characterization of the following colony traits: form, elevation, margins, opacity, surface, iridescence, texture, shine, and color. Distinct colonies were streaked to purity on MA/2 plates and then transferred to liquid Marine Broth (Difco™) cultures, which were incubated for 48-72h at 20-22ºC in continuous movement. This process was repeated to ensure complete isolation of the bacterial symbionts.

### 2.3 Cryopreservation of the bacterial isolates

Pure cultures were cryopreserved using glycerol as a cryoprotectant (Pegg, 2007). For this, we mixed in sterile cryovials (DeltaLab) 0.5 mL of liquid culture and 0.5 mL of 50% glycerol (diluted in distilled water). The cryovials were stored at -80ºC.

### 2.4 DNA extraction and 16S rRNA sequencing

DNA was extracted from pure cultures using the Quick-DNA Microprep Kit (Zymo Research). To initiate the extraction protocol, colonies from each isolate were transferred to sterile Eppendorf ® tubes containing 400 μL of lysis buffer from the kit. After cell lysis, the next steps indicated by the manufacturer were followed. The extracted DNA was stored at -20ºC until further processing. The 16S rRNA gene was amplified using the universal bacterial primers 337F (5’ GACTCCTACGGGAGGCWGCAG 3’) and 907 RM (5’ CCGTCAATTCMTTTGAGTTT 3’). The PCR mixtures (25 μL) contained 1.25 μL of each primer, 2 μL of DNA template, 12.5 μL of PCR Master Mix (Invitrogen by Thermo Fisher Scientific) and 8 μL of nuclease-free water. A thermocycler was used to perform the PCR reaction, programmed to run as follows: 5 mins at 94ºC, 30 cycles of 94ºC (5 mins) – 55ºC (1 min) – 72ºC (2 mins), and 10 mins at 72ºC for the final extension. A gel electrophoresis (1% agarose gel) confirmed the correct amplification of the gene. Correctly amplified samples were sequenced by Macrogen (https://macrogenclinical.com/) using Sanger sequencing.

### 2.5 Identification and phylogenetic analyses

Raw sequencing data for this project has been deposited on NCBI under the accession numbers XXX-XXX. The DNA sequence chromatograms were visualized and cleaned using Chromas 2.6.6. Clean sequences were subjected to a BLAST search against the latest release of GenBank for taxonomic identification. Sequences assigned to terrestrial or human-associated bacteria were considered potential contaminants and excluded from subsequent analysis. Sequences were clustered at 97% identity using vsearch 2.23.0, assuming that bacteria with >97% sequence identity represent individual operational taxonomic units or OTUs (Rognes et al., 2016; Stackebrandt & Goebel, 1994).

The 16S rRNA sequences from the isolates, along with sequences from members of related genera (downloaded from the NCBI database), were aligned using MAFFT v7.505, and the alignment was inspected in AliView v1.28 (Katoh & Standley, 2013; Larsson, 2014). Then, trimAI v1.4 was employed to clean the alignment using automated trimming (Capella-Gutiérrez et al., 2009). Phylogenetic trees were inferred through maximum-likelihood using RAxML v8.2.12 (Stamatakis, 2006). The resultant trees were visualized and edited using the Interactive Tree of Life (iTOL) online platform (Letunic & Bork, 2007).

## 3. RESULTS

A total of 134 bacterial colonies were selected for isolation after the initial morphological characterization. After the complete isolation protocol, 96 bacterial isolates were obtained. This reduction in the final number of isolates resulted from the removal of morphologically redundant colonies and the inability of some bacteria to grow in specific media at different stages. Most of these 96 bacteria were cultured in MA/2 medium; only 12 isolates were obtained from MA/2 supplemented with coral extract. Although not many isolates were obtained from this medium, it is worth noting that 2 of them were exclusively growing in this medium (“Cultured bacterium 4J Esin”, from the genus *Enterovibrio*, and “Cultured bacterium 4G Esin”, from the genus *Pseudoaltermonas*). In the case of TCBS Agar, no growth was observed. Longer incubation times (up to two and three weeks) resulted in the isolation of slower-growing genera such as *Leisingera, Roseibium, Tritonibacter*, and *Vibrio* from the clade Halioticoli from MA/2. Replicates from the 96 isolates were stored in the cryobank. After cryopreservation, cultures were regrown to verify their viability following freezing.

The 16S rRNA gene of these 96 isolates was sequenced, and a BLAST search identified 12 contaminants (i.e., isolated bacteria that were not coral symbionts). Therefore, the presented results are based on a total of 84 isolates identified as coral-associates. Colony morphology was used to distinguish different strains. For each colony, a total of 8 different traits were examined (elevation, form, iridescence, margin, opacity, shine, surface and texture), but often only one of the categories dominated per trait (Figure 2). For example, most colonies were predominantly circular, with entire margins, raised in elevation, and with a smooth surface and moist texture. In contrast, other traits, such as mucoid texture or a rough surface, were rare.

**Figure 2.**
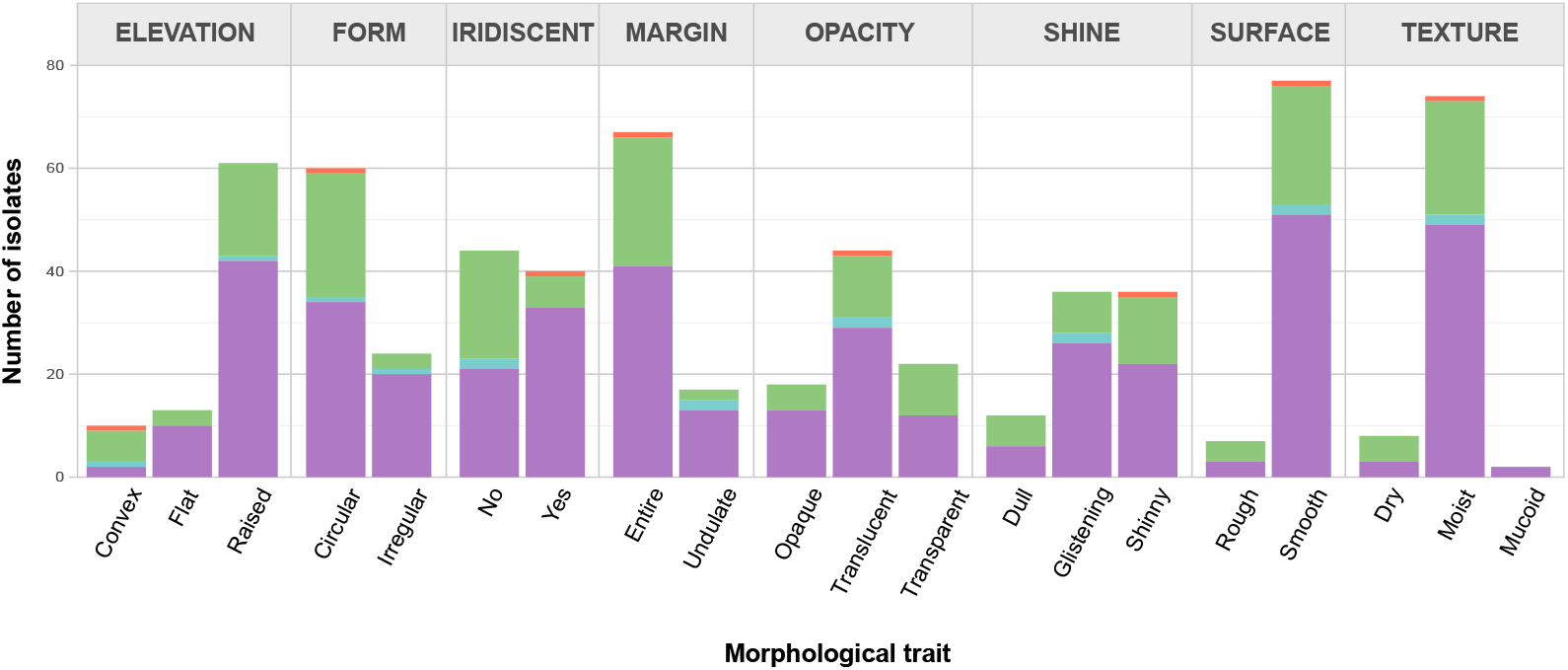
Morphological characterization of the 84 bacterial isolates. Bar plot showing the distribution of isolates across eight colony traits (elevation, form, iridescence, margin, opacity, shine, surface, and texture). Bars represent the number of isolates per subtrait and are color-coded by bacterial class.

The different strains were grouped into Operational Taxonomic Units (OTUs) by clustering their sequences at 97% similarity. The assessment of the phenotypic characteristics helped reduce redundancy in the isolation process, with only 32% of the OTUs (12 out of the 37) being isolated more than once (Figure 3). Those OTUs that were isolated more than once exhibit morphological variability in 83% of the cases, which explains their repeated isolation. However, human error led to the repeated isolation of 2 OTUs despite having identical colony morphologies (Figure S1).

**Figure 3.**
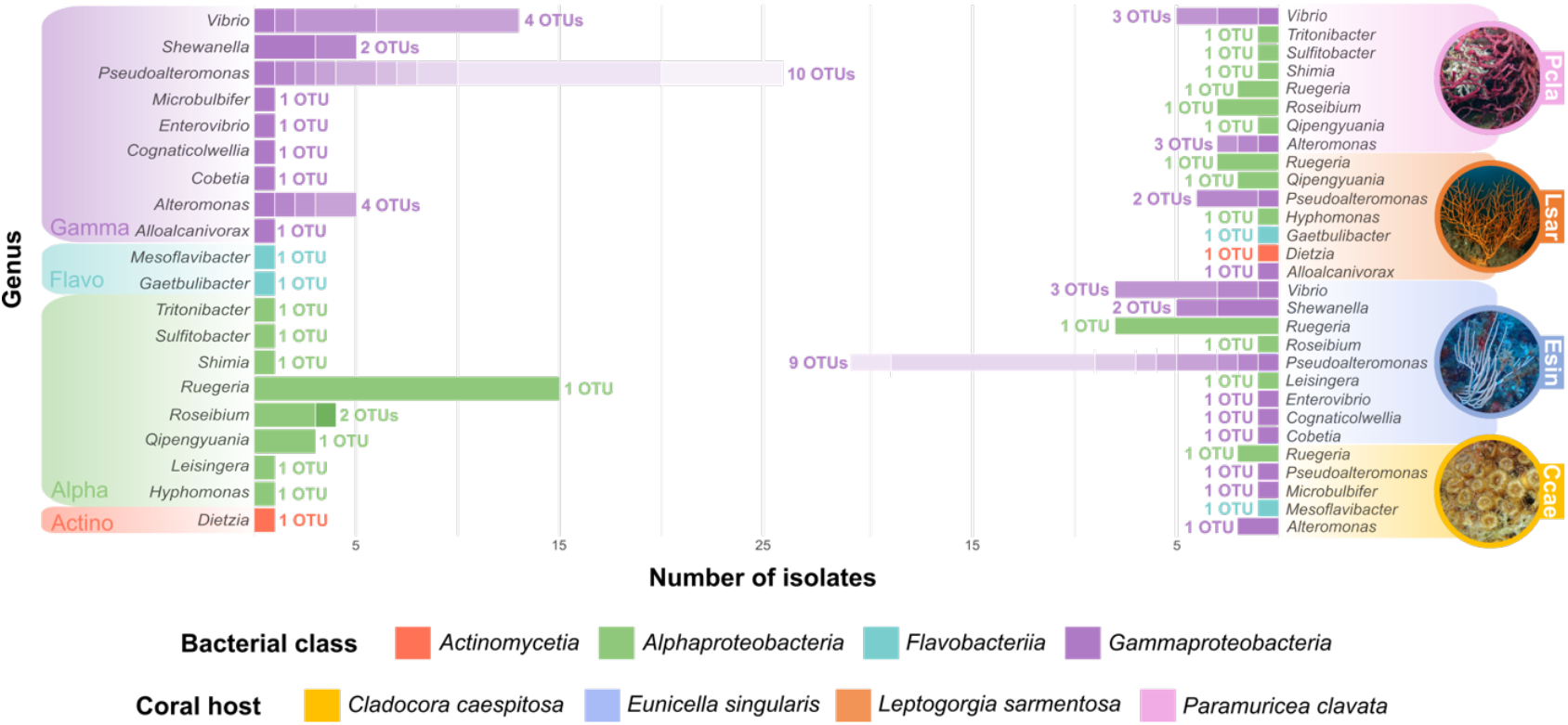
Taxonomic diversity and host distribution of cultured coral-associated bacteria. Left panel: number of isolates per genus (20 genera across four bacterial classes). Right panel: host-specific distribution of these bacterial genera across four coral species: *C. caespitosa, E. singularis, L. sarmentosa*, and *P. clavata*. The shading changes within bars indicate distinct OTUs (97% similarity).

In total, 37 different OTUs were successfully isolated, which we classified into 4 classes (Alphaproteobacteria, Gammaproteobacteria, Flavobacteriia and Actinobacteria) and 20 genera (Figure 3). Alphaproteobacteria and Gammaproteobacteria isolates showed greater colony variability. However, we can observe certain tendencies such as the prevalence of iridescence in Gammaproteobacteria or the clear dominance of entire and circular colonies in Alphaproteobacteria. The morphologies of the two isolates from Flavobacteriia are similar in all traits except elevation (raised or convex) and form (circular or irregular).

Regarding the diversity of the isolates (Figure 3), most of the genera are represented by a single OTU. Conversely, genera like *Vibrio* (4 OTUs), *Alteromonas* (4 OTUs) and especially *Pseudoalteromonas* (10 OTUs) stand out for their greater strain diversity. Both, a repeated isolation (such as in *Ruegeria*) and a high strain diversity (as in *Vibrio* and *Pseudoalteromonas*) contribute to the abundance of certain genera which surpass the average number of isolates per genus.

The final number of isolates and OTUs varies greatly among coral hosts (Figure 3). *E. singularis* was the coral from which more bacteria were retrieved: a total of 47 isolates encompassing 20 different OTUs. At the opposite end, from *C. caespitosa*, only 7 bacteria were isolated, comprising 5 OTUs. From *P. clavata*, 17 isolates were obtained and divided into 12 distinct OTUs. Finally, from *L. sarmentosa*, 13 bacteria belonging to 8 OTUs were isolated. Those genera with greater diversity in *E. singularis* were *Pseudoalteromonas* (9 OTUs) and *Vibrio* (3 OTUs). In comparison, in *P. clavata*, the genera *Vibrio* and *Alteromonas* were the most diverse (both with 3 OTUs). In *L. sarmentosa*, the genus *Pseudoalteromonas* was the only genus with more than one strain (2 OTUs), whereas in *Cladocora* each genus was represented by a single OTU.

At the genus level, there are differences in the bacterial microbiota composition across corals (Figure 4). Only the genus *Ruegeria* was present in all corals. Bacteria from the genus *Pseudoalteromonas* were isolated from three hosts (*C. caespitosa, E. singularis*, and *L. sarmentosa*), but no species from this genus were isolated from *P. clavata*. Furthermore, various genera were found exclusively in every host: 2 in *C. caespitosa*, 5 in *E. singularis*, 4 in *L. sarmentosa*, and 3 in *P. clavata*. In total, 14 out of 20 isolated genera (70%) were specific to a single coral species. Notably, even the genera shared among corals comprised host-specific strains, with only one OTU from the genus *Ruegeria* common to all analyzed corals. In other words, only 2.7% of the isolated bacteria were shared by all hosts, and 86.5% were specific to individual coral species (Figure S2).

**Figure 4.**
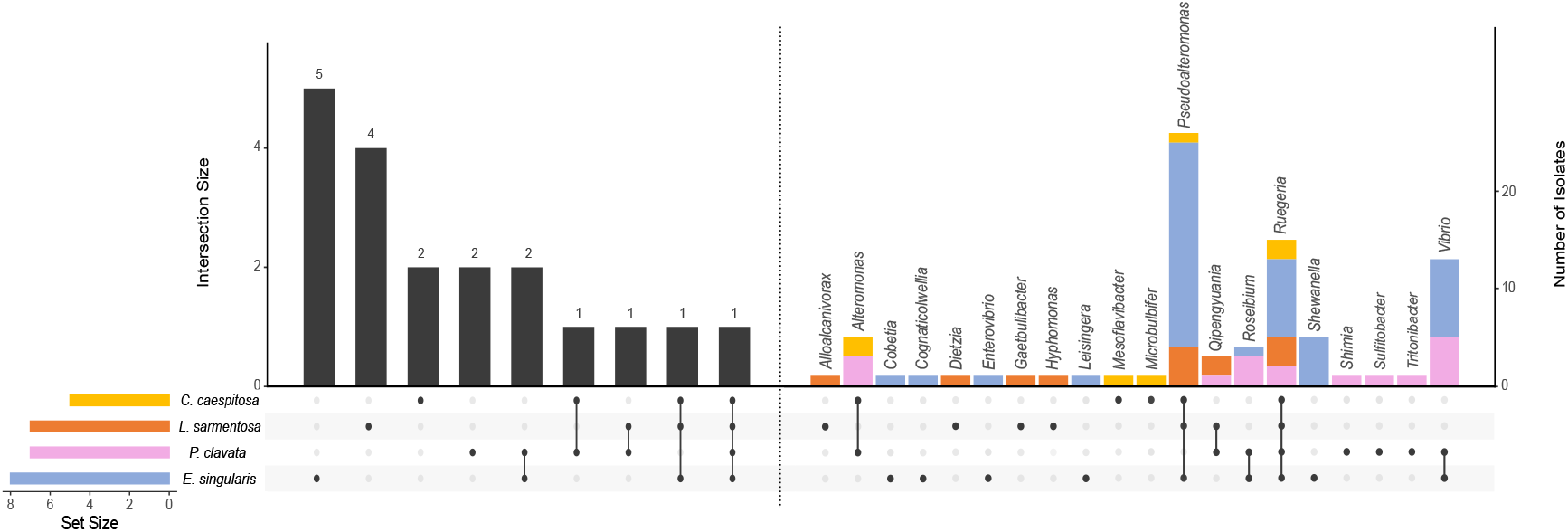
Shared and host-specific bacterial genera among coral hosts. UpSet plot (left) showing shared and exclusive genera among the four coral species, and bar plot (right) providing taxonomic information for each intersection, displaying isolate numbers per genus, color-coded by host. Thirteen genera were unique to a single host, whereas *Ruegeria* occurred in all microbiotas.

Seven phylogenetic trees were generated, each corresponding to specific bacterial groups of interest. Within the class Gammaproteobacteria, which was the most abundant one, we generated three different trees: one for Alteromonadales (Figure 5), one that zooms into the family *Vibrionaceae* (Figure 6), and one that combines Oceanospirillales and Cellvibrionales (Figure S3). For the class Alphaproteobacteria, two separate phylogenies were generated: one including *Roseobacteraceae, Erythrobacteraceae*, and *Hyphomonadaceae* (Figure S4), and a second for *Stappiaceae* (Figure S5). Additionally, single phylogenetic trees were produced for the remaining classes - one for Actinobacteria (family *Dietziaceae*) (Figure S6) and another for Flavobacteriia (family *Flavobacteriaceae*) (Figure S7).

**Figure 5.**
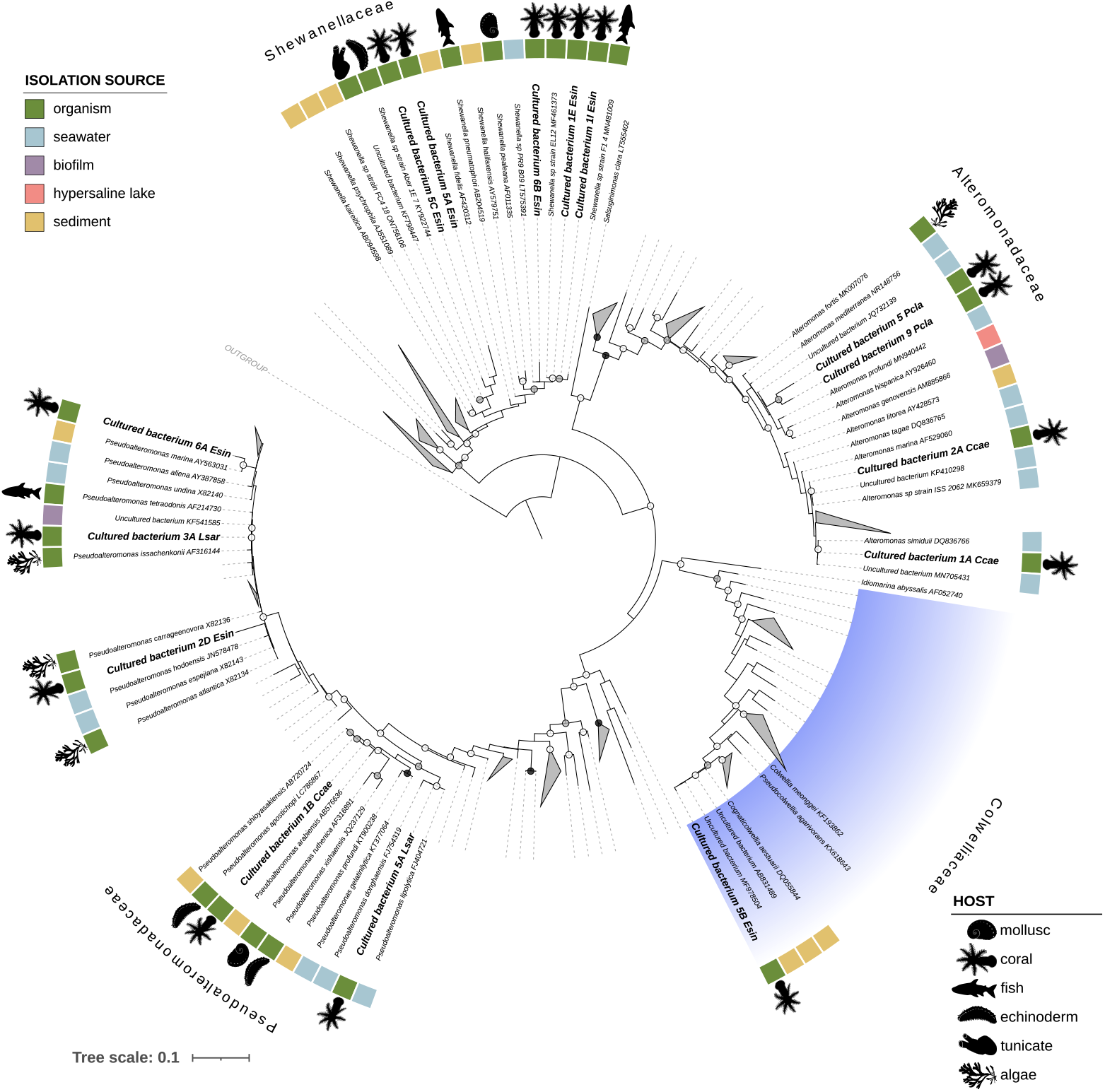
Maximum-likelihood phylogeny of Alteromonadales based on 16S rRNA gene sequences. The tree includes isolates obtained in this study alongside closely related reference sequences; Pseudomonadales were used as outgroup. Bootstrap support (1000 replicates) is indicated by circles at nodes (black, 100%; grey, 90–99%; white, 70–89%). Major clades are color-coded by family (Alteromonadaceae, Colwelliaceae, Pseudoalteromonadaceae, and Shewanellaceae). Colored squares at the tips indicate isolation source, and animal silhouettes inform about the host of origin for those strains detected in organisms.

**Figure 6.**
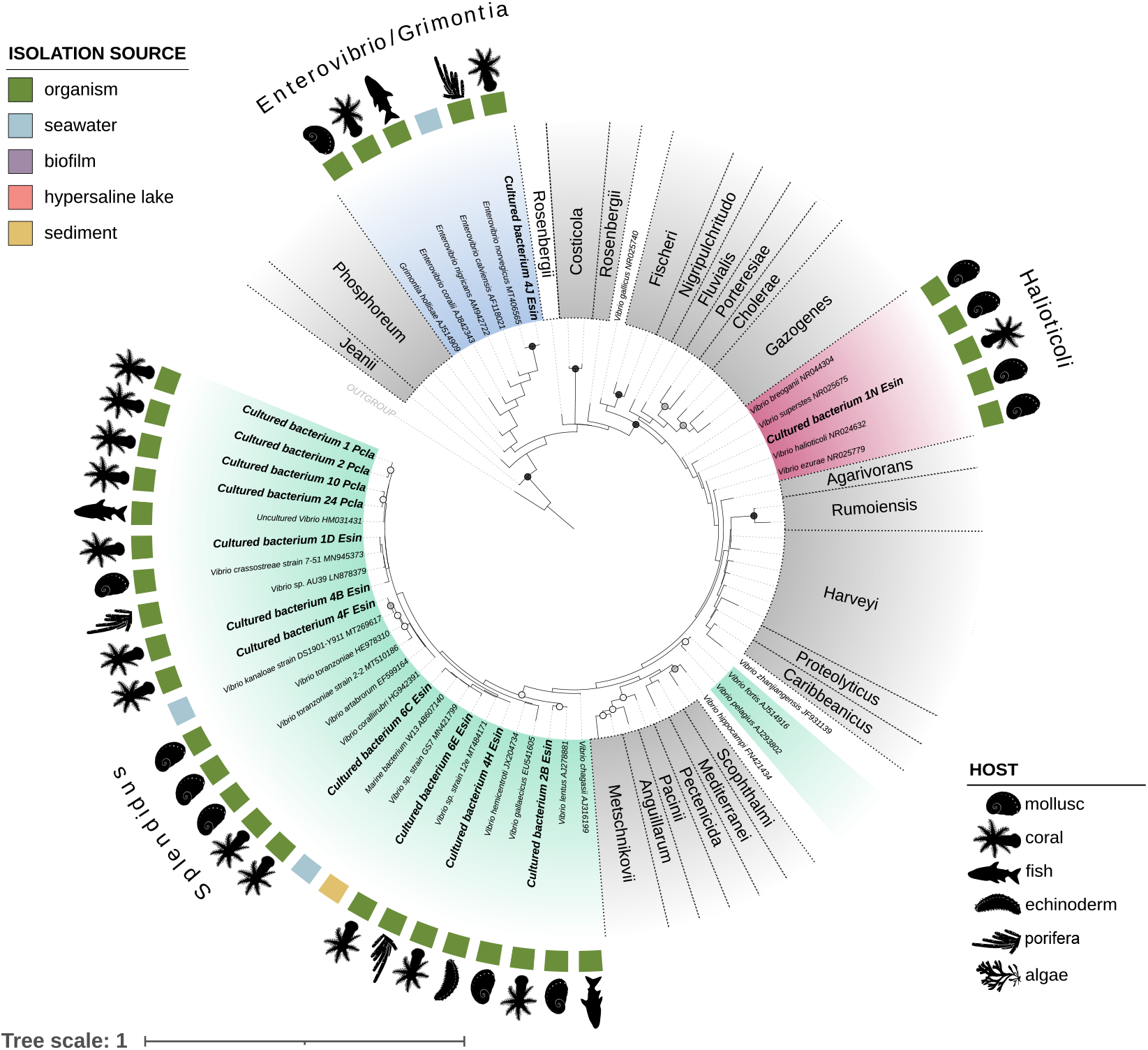
Maximum-likelihood phylogeny of Vibrionaceae based on 16S rRNA gene sequences. The tree includes isolates obtained in this study alongside closely related reference sequences; Shewanellaceae were used as outgroup. Bootstrap support (1000 replicates) is indicated by circles at nodes (black, 100%; grey, 90–99%; white, 70–89%). The three clades of interest (Splendidus, Halioticoli, Enterovibrio/Grimontia) are color-coded, while for the remaining ones only the clade name is provided. Colored squares at the tips indicate isolation source, and animal silhouettes inform about the host of origin for those strains detected in organisms.

The phylogenetic analysis of the order Alteromonadales (Figure 5) includes isolates’ sequences from the families *Shewanellaceae, Alteromonadaceae, Colwelliaceae* and *Pseudoalteromonadaceae*. These isolates were obtained from *E. singularis* (8 sequences), *L. sarmentosa* (2 sequences), *P. clavata* (2 sequences) and *C. caespitosa* (3 sequences). The resulting phylogenetic tree comprises 212 sequences, 15 of which are from our bacterial isolates. From the family *Shewanellaceae*, a total of 5 bacteria were isolated (“5C Esin”, “5A Esin”, “6B Esin”, “1E Esin” and “1I Esin”), all of them associated within the host *E. singularis*. The two remaining Alteromonadales species isolated from this coral belonged to the families *Colwelliaceae* (“5B Esin”) and *Pseudoalteromonadaceae* (“2D Esin”). Focusing on *P. clavata*, both isolates belonged to the family *Alteromonadaceae* (“5 Pcla” and “9 Pcla”), and the same pattern was observed in *L. sarmentosa*, as the 2 isolates belonged to the family *Pseudoalteromonadaceae* (“5A Lsar” and “3A Lsar”). In contrast, the bacteria isolated from *C. caespitosa* include members of both families: two *Alteromonadaceae* (“2A Ccae” and “1A Ccae”) and one *Pseudoalteromonadaceae* (“1B Ccae”).

The phylogenetic analysis of the family *Vibrionaceae* (Figure 6), includes the 16S rRNA gene sequences obtained from *Vibrio* and *Enterovibrio* isolates. The isolates come from the corals *E. singularis* (9 sequences) and *P. clavata* (4 sequences), as no other bacteria from the family *Vibrionaceae* were isolated from *C. caespitosa* nor *L. sarmentosa*. The resultant phylogenetic tree includes 80 sequences, 67 of which provide phylogenetic context, and 13 originate from the present project. The phylogenetic tree reveals that the isolated bacterial symbionts belong to 3 different *Vibrionaceae* clades: Halioticoli clade (“1N Esin”), Splendidus clade (“1 Pcla”, “2 Pcla”, “10 Pcla”, “24 Pcla”, “1D Esin”, “4B Esin”, “4F Esin”, “6E Esin”, “6C Esin”, “4H Esin”, “2B Esin”) and the super-clade Salinivibrio-Grimonita-Enterovibrio (“4J Esin”). The coral *E. singularis* presented representatives from the three different clades, while *P. clavata* vibrios only belong to the clade Splendidus.

## 4. DISCUSSION

### 4.1 Culturing the coral-associated bacterial diversity

In this study, using a culture-based approach, we were able to explore the diversity of bacterial symbionts associated with healthy colonies of *Cladocora caespitosa, Leptogorgia sarmentosa, Eunicella singularis*, and *Paramuricea clavata*. We successfully isolated and taxonomically identified 84 coral-associated bacterial strains, recovering a diverse fraction of the bacterial microbiota (37 OTUs) and highlighting specific community patterns per coral host. Previous studies have predominantly relied on 16S rRNA amplicon sequencing or focused on specific taxa of interest, such as putative pathogenic agents (e.g., *Vibrio* spp.) (Grottoli et al., 2018; Rubio-Portillo et al., 2018a). Nevertheless, complementing culture-independent studies by isolating pure cultures of bacterial symbionts is essential for linking taxonomic diversity to symbiont morphological, functional, and genomic traits (Li et al., 2022; Sweet et al., 2021).

Most isolates were obtained from the coral E. singularis (47 isolates), which may be related to the larger number of initial coral samples used (five individuals). Conversely, using a unique sample per species may have accounted for the lower number of isolates from the other three corals. However, it is also plausible that the higher number of isolates obtained from *Eunicella* is explained by this species having an easily culturable prokaryotic microbiota (Keller-Costa et al., 2017).

Culture-based approaches to studying the microbiome will inherently reveal only a subset of the bacterial symbionts associated with corals due to the so-called “great plate count anomaly” (Schultz et al., 2022). This phenomenon refers to the difficulty (or impossibility) of culturing the entire existing bacterial diversity under laboratory conditions. However, our results demonstrate that the cultivable fraction recovered through culture-based studies qualitatively reflects dominant bacterial groups frequently detected through molecular techniques (Huggett & Apprill, 2019; Rohwer et al., 2002; Sweet et al., 2021). This suggests that, despite its initial limitations, culturing remains a useful tool for the in-depth study of ecologically relevant members of the coral microbiome. Indeed, previous studies have shown that a taxonomically diverse array of bacteria can be isolated using various media and culture conditions (Schultz et al., 2022; Sweet et al., 2021).

We should interpret the “great plate count anomaly” as a limitation of current methodology rather than as an intrinsic inability of bacteria to grow in vitro. Classical microbiological techniques are limited and thus may fail to recreate suitable environments for different bacterial groups to thrive, shifting the focus from “unculturable” to “yet-uncultured” bacteria (Schultz et al., 2022; Sweet et al., 2021). In this study, the use of three different media was intended to overcome some of these potential constraints and support the culture of a diverse bacterial community. Among them, the media that worked the best was MA/2, consistent with previous studies (Sweet et al., 2021). The dilution of this nutrient-rich agar was key to allowing the growth of slower-growing taxa that would otherwise be outcompeted by fast growers (Sipkema et al., 2011). In contrast, no growth was obtained on TCBS agar despite the presence of *Vibrio* sp. among our isolates. It is possible that highly selective media might be useful for recovering free-living taxa but not coral-associated bacteria. On the other hand, adding coral extract to MA/2 enabled the isolation of 12 bacterial strains, suggesting that certain host-derived compounds may be crucial for the growth of specialized symbionts (Schultz et al., 2022). In addition, longer incubation times (of up to three weeks) allowed the recovery of slower-growing species that could be easily overlooked (Keller-Costa et al., 2017; Pulschen et al., 2017). Overall, these results highlight the importance of diversifying classical culturing procedures when trying to culture unknown coral-associated microorganisms (Lewis et al., 2021).

### 4.2 The importance of assessing colony morphology

Morphological characterization of bacterial colonies proved a useful preliminary tool for selection to avoid repeated isolation. However, it is an imperfect proxy for taxonomic identity, owing to both intrinsic variability within the same OTU and human errors. Therefore, we recommend using morphological characterization as a starting point, always combined with molecular identification.

In several cases, we observed different colony morphologies for the same OTU. For instance, one *Pseudoalteromonas* OTU exhibited two phenotypes that differed in texture and shine: one being mucoid and glistening, and the second one being moist and shiny. According to the literature, some colony characteristics can vary with external factors such as nutrient availability (Martinez-Garcia et al., 2022). When examining different OTU morphologies across culture conditions, we identified the following scenarios: (a) different media resulting in different morphologies for the OTU (e.g. OTU 23), (b) different media but identical OTU morphology (e.g. OTU 20), and (c) distinct morphologies arising even in the same medium (e.g. OTU 36). Therefore, while variation in culture media may explain some morphological differences (Martinez-Garcia et al., 2022), the occurrence of different morphologies under identical conditions suggests that there are additional factors involved. For example, phenotypic heterogeneity (defined as phenotypic diversity among genetically identical individuals living under the same conditions) has been widely reported as an adaptive strategy in marine bacterial populations (Ackermann, 2015). This phenomenon allows bacteria to maintain functional diversity and persist in fluctuating environments (Ackermann, 2015), what may be important for establishing long-term symbiotic associations and survive as part of the coral holobiont. This plasticity has important methodological implications, as it may lead to repeated isolation of the same species if they develop different phenotypes or, conversely, to the erroneous assumption of identical taxa when distinct species display similar morphologies under the same conditions.

Despite these limitations, continued assessment of colony morphology remains valuable in microbiological studies, as it can provide insights into ecological processes. For example, “sticky” variants (here classified as mucoid) have been associated with hyperadherence, autoaggregation, and enhanced biofilm-forming ability (Kirisits et al., 2005), traits that could facilitate bacterial association with the coral host. In this context, the uniformity observed for certain traits in our isolates suggests that colony morphology might reflect biologically meaningful adaptations rather than accidental variation. Interestingly, the only strain shared by all hosts, belonging to the genus *Ruegeria*, exhibited rough surfaces and dull, dry colonies. Previous studies have linked rough or rugose colonies to increased exopolysaccharide production, which has been described as necessary for adherence and biofilm development in aquatic environments (Kirisits et al., 2005; Watnick et al., 2001). Notably, several colonies (especially Gammaproteobacteria – Table S1) displayed iridescent phenotypes, a structural coloration often resultant from a high cell organization. This cellular organization has been linked to coordinated behaviors such as biofilm formation, quorum sensing and surface motility, processes that potentially facilitate surface attachment and stable host-associated biofilms (Zomer et al., 2024). Particularly, gliding motility, which was widely exhibited by our isolates, has been linked with iridescence and successful surface colonization (Kientz et al., 2012; Zomer et al., 2024). Furthermore, both gliding motility and iridescence, have also been associated with the ability to degrade complex polysaccharides (Gavriilidou et al., 2020; Zomer et al., 2024), suggesting that these bacteria might play a role in degrading polymers found in the coral mucus. Supporting this, growth of some colonies caused weakening of the Marine Agar surface, consistent as well with enzymatic degradation of polymers by agarolytic bacteria (Chi et al., 2012). Together, these findings suggest that colony morphology may reflect traits relevant for host association and symbiosis establishment, while also providing valuable insights into possible ecological roles of bacteria within the holobiont. However, more research is needed regarding which phenotypic characteristics to enable bacteria to associate with corals.

### 4.3 Culturable bacterial diversity associated with Mediterranean corals

As noted previously, culture-based studies recover only a subset of the total bacterial diversity associated with coral hosts. Nevertheless, our results qualitatively align with those of previous studies. Most of the bacteria isolated belonged to the phylum Proteobacteria, specifically to the classes Gammaproteobacteria and Alphaproteobacteria. Similar culture-based studies of the coral microbiota have repetitively found a high abundance of these two bacterial classes across different hosts and regions (Sweet et al., 2021). Although less abundant, members of Flavobacteriia and Actinomycetia have also been described in previous works as coral-associated bacteria (Sweet et al., 2021). Concerning molecular-based studies, the community is dominated by four phyla: Proteobacteria, Bacteroidetes, Actinobacteria, and Firmicutes (Huggett & Apprill, 2019; Rohwer et al., 2002; Sweet et al., 2021), having isolated in this study representants from three of these phyla. Thus, our results are consistent with previous culture-based and molecular-based studies. At the genus level, the most common isolates have also been described before as typical components of the coral holobiont (Sweet et al., 2021). However, no bacteria from the genus Endozoicomonas, previously described as a principal component of the bacterial microbiota, were isolated in this work, reflecting the traditional difficulty of culturing this genus in vitro (Keller-Costa et al., 2021; van Oppen & Blackall, 2019).

Our results also underscore a marked host specificity of the bacterial microbiota. This pattern is consistent with previous studies indicating that coral species largely shape the composition of their associated prokaryotic communities, even across different geographic regions (Littman et al., 2009; Morrow et al., 2012). The isolates obtained in this study were predominantly associated with a single coral host. Indeed, among all of them, only *Ruegeria* is shared by all corals, a genus that has been documented before to be highly abundant across different coral samples and part of the core microbiome (Huggett & Apprill, 2019; Rubio-Portillo et al., 2021). Interestingly, even the different *Ruegeria* isolates shared more than 97% 16S rRNA gene sequence similarity, therefore constituting a single OTU, consistent with patterns previously reported for Mediterranean corals (Rubio-Portillo et al., 2021).

#### 4.3.1 Phylogenetic insights into dominant bacterial groups

Phylogenetic analyses provide the necessary evolutionary context to interpret the ecological relevance and potential novelty of the bacterial isolates recovered in this study. The Alteromonadales and Vibrionaceae trees reflect the evolutionary relationships between the isolates and closely related species, placing them into well-defined families (*Alteromonadaceae, Colwelliaceae, Pseudoalteromonadaceae* and *Shewanellaceae*) and clades (Splendidus, Halioticoli and Enterovibrio/Grimontia superclade), respectively. Across both analyses presented here, many of the isolates cluster with other sequences recovered from marine organisms, which further supports the identity of the isolated bacteria as members of the coral microbiome.

Within Alteromonadales, the analysis revealed that many of our isolates - especially those belonging to the families Shewanellaceae and Pseudoalteromonadaceae - are closely related to other bacteria that associate with marine organisms. Indeed, we even find a bacterial isolate sister to the isolate “6B Esin” which was isolated from the gorgonian species *Eunicella labiata* (Keller-Costa et al., 2017). Other relatives could be retrieved from diverse marine organisms, including fish, algae, tunicates, echinoderms, and mollusks (Figure 5). In contrast, the isolate from Colwelliaceae clustered with species previously detected in deep-sea and tidal flat sediments. Nevertheless, members of Colwelliaceae have also been reported in the microbiomes of other cnidarians (Khabibulina et al., 2025) and have been associated with growth-promoting effects in marine organisms, including algae (Liu et al., 2023). Regarding the Vibrionaceae tree, we find an even higher level of relatedness with lineages that associate with marine organisms (Figure 6). Most of the obtained isolates fell within the Splendidus clade, in which relatives are isolated from diverse organisms including mollusks, sponges, corals, echinoderms, and fish (Esteves et al., 2016; Lasa et al., 2013). In contrast, a single bacterium from the Halioticoli clade was isolated; this clade primarily comprises mollusc-associated bacteria. Additionally, the Enterovibrio isolate grouped within the SGE superclade (Salinivibrio-Grimontia-Enterovibrio) and its 16S sequence was identical to the species *E. norvegicus*, which associated with the sponge *Halichondria panicea* (Schmittmann et al., 2022).

The placement of certain isolates in these phylogenetic trees suggests that they represent previously uncharacterized bacterial species. Within Alteromonadales, all *Shewanella* isolates and several *Alteromonas* (“5 Pcla”, “9 Pcla”, “2A Ccae”) clustered with environmental sequences or uncultured strains rather than with formally described species. In relation to *Shewanella* isolates, the closest related species were also isolated from marine organisms including corals, sea urchins, fishes and ascidians (Dishaw et al., 2014; Keller-Costa et al., 2017; Laport et al., 2018; Tesdorpf et al., 2022). Similarly, multiple *Vibrio* isolates formed well-defined groups distinct from known species. Within the Splendidus clade, we can distinguish two unresolved lineages that encompass putative novel taxa isolated in this study. One lineage comprised the isolates obtained from *P. clavata* (“1 Pcla”, “2 Pcla”, “10 Pcla” and “24 Pcla”), and its closest relative was detected through eDNA in fish guts. The second clade included three isolates derived from *E. singularis* (“6C Esin”, “6E Esin”, “4H Esin”), and was most closely related to species recovered from sponges and environmental samples (Rodriguez Jimenez et al., 2021). While previous studies detected members of these lineages using culture-independent methods, this study isolated these previously uncultured organisms. These results underscore the importance of studying through cultures the microbial communities associated with corals, as they can harbor a substantial hidden bacterial diversity that may be essential for holobiont functioning.

Additional phylogenetic analyses of the remaining isolates are provided as supplementary figures (Figures S3-S7). These analyses highlight the broad taxonomic diversity encompassed within the culturable fraction of the coral microbiome, and further support the patterns described for Alteromonadales and Vibrionaceae. Most of the isolates cluster with sequences previously retrieved from ocean habitats or marine species, including other corals as well as diverse hosts like sponges, ciliates, cyanobacteria, fish and crustaceans. Moreover, these groups likewise contain several lineages that branch apart from formally described species (e.g. “13 Pcla”, “22 Pcla”, and “23 Pcla” within Roseobacteraceae, or “2C Ccae” within Flavobacteriaceae), suggesting the presence of additional potentially novel taxa within these families.

#### 4.3.2 Potential ecological roles within the coral holobiont

Coral-associated bacteria have been recognized as key members of the holobiont, performing a wide variety of roles that contribute to its functioning (Keller-Costa et al., 2021; Rosenberg et al., 2007; Webster & Reusch, 2017). However, the specific roles of bacterial symbionts are usually difficult to assess, especially if the only available information comes mostly from barcode gene sequences. Based on previously available data, we could identify putative functions of the isolates such as with carbon fixation, N and S cycling, host defense, and metabolite synthesis (complex carbohydrates, vitamins, amino acids…).

For instance, the bacterial community associated with corals contained carbon-fixing bacteria such as *Ruegeria* (via the Calvin cycle), *Leisingera, Erythrobacter*, and *Sulfitobacter* (viaanoxygenic photosynthesis) (Li et al., 2022). In nutrient cycling, genera including *Pseudoalteromonas* and *Sulfitobacter* have been associated with the oxidation of nitrite to nitrate, whereas *Ruegeria* and members of Alteromonadales may participate in the degradation of DMSP (Li et al., 2022; Raimundo et al., 2018). Furthermore, several isolates may also contribute to the defense against pathogens. Genera such *Pseudoalteromonas, Ruegeria*, or *Cobetia* can protect corals through production of antimicrobial compounds or resource competition (Rosado et al., 2019; Rubio-Portillo et al., 2021; Sweet et al., 2021).

Notably, the genus *Ruegeria* has been proposed as a probiotic for corals since it inhibits the growth of other potentially pathogenic bacteria, such as *V. coralliilyticus*, and modulates coral-associated bacterial community (Kitamura et al., 2021; Rubio-Portillo et al., 2021; Xu et al., 2024). The consistent recovery of this genus across the four coral hosts studied aligns with previous studies identifying it as a core microbiome member (Rubio-Portillo et al., 2021). Although previously associated with the global spread of coral diseases (Apprill et al., 2013; Sekar et al., 2008; Séré et al., 2013), its stable presence across coral species, conditions, and regions supports an important role in coral holobiont functioning.

In contrast, other genera such as *Vibrio* have been repeatedly reported to be linked with coral diseases, as their abundance is usually higher in necrosed tissues (Bisanti et al., 2025; Munn, 2015; Rubio-Portillo et al., 2021; Vezzulli et al., 2010). Pathogenicity assays pinpoint *V. coralliilyticus* as a causative agent of coral disease (Ushijima et al., 2020). However, Vibrio species have also been detected in the microbiomes of healthy corals and other marine organisms (Bisanti et al., 2025; Rubio-Portillo et al., 2021; Rubio-Portillo et al., 2018b). This indicates that their specific ecological role might be context-dependent, potentially shifting opportunistically from a mutualistic to a pathogenic lifestyle under certain conditions, such as thermal stress or dysbiosis. For instance, the co-occurrence of multiple vibrio species (such as *V. coralliilyticus* and *V. mediterranei*) has been shown to enhance virulence, suggesting that species interactions can also be determinant for the development of disease (Rubio-Portillo et al., 2018b; Ushijima et al., 2020).

Further functional analysis and in vitro experimentation are needed to elucidate the complex interactions that occur within the host. In this context, the current generation of cultures represents a crucial step towards the comprehensive understanding of the coral microbiota.

## 5. CONCLUSION

In the present study, we successfully isolated 84 pure cultures of symbiotic bacteria from four Mediterranean corals (*C. caespitosa, L. sarmentosa, E. singularis*, and *P. clavata*). Using a culture-based approach, we recovered a substantial diversity of coral-associated bacteria, which taxonomically reflects the dominant bacterial groups and community structure reported in previous molecular studies. Phylogenetic analyses revealed the close relatedness between our isolates and other host-associated marine bacteria. In addition, several isolates clustered with uncharacterized bacteria, indicating that the coral microbiome harbors unexplored microbial diversity.

Overall, these results emphasize the importance of integrating current holobiont information, obtained mainly through molecular techniques, with microbial culturing to achieve a more comprehensive understanding of coral-associated bacterial diversity and function, and coral-microbe interactions. The culture cryobank generated here provides a foundation for future genomic and experimental studies, which are essential for elucidating the roles of bacteria within the coral holobiont and their relevance to coral health and resilience.

## AUTHOR CONTRIBUTIONS

**Rocío Mozo:** investigation, methodology, data curation, formal analysis, visualization, writing – original draft. **Andrea Illa-Oviedo:** investigation, methodology, writing – review and editing. **Javier del Campo:** conceptualization, methodology, funding acquisition, project administration, resources, supervision, validation, writing – review and editing.

## ACKNOWLEDGMENTS

We thank Janire Salazar and Diego Kersting for their collaboration with field sampling, the coral samples they provided were essential for the development of the study. We are also grateful to Guillem Manau for providing the coral photographs used for the figures of this study. Computational analyses were supported by the high-performance computing services of CESGA (Centro de Supercomputación de Galicia), whose resources and technical assistance were essential for this work.

## DATA AVAILABILITY STATEMENT

The 16S rRNA gene amplicon sequencing data will be deposited on NCBI under the accession numbers XXX-XXX. The raw .tre files generated for the phylogenetic analyses are available upon request.

