## Supplementary Figures for "CULTURE AND ISOLATION OF BACTERIA ASSOCIATED WITH MEDITERRANEAN CORALS"

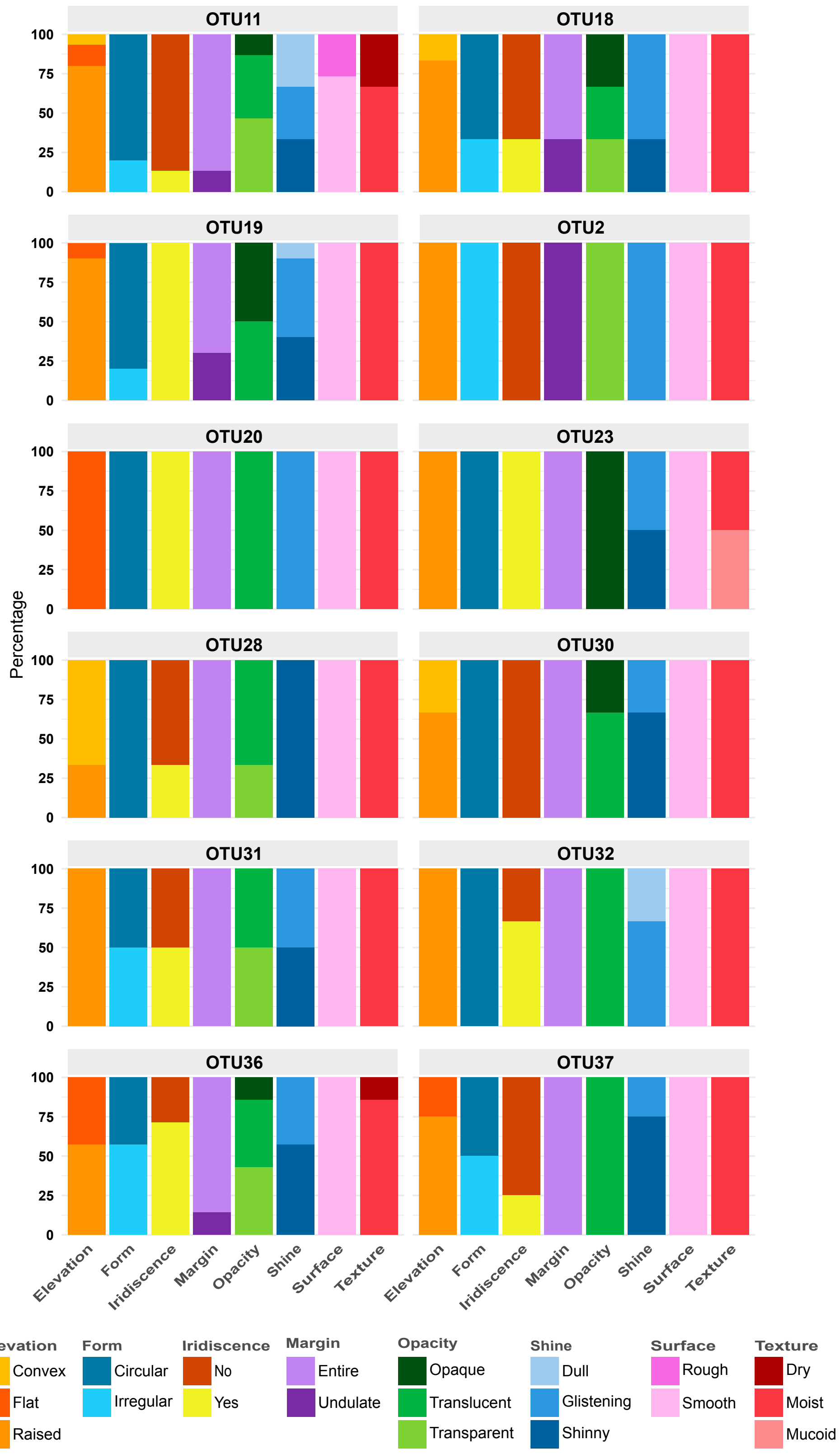


**Supplementary Figure 1. Morphological variation of repeatedly isolated OTUs.** Relative percentage distribution of colony traits for the 12 OTUs isolated multiple times. Most OTUs exhibited variation in at least one trait, whereas others (e.g. OTU2 and OTU20) displayed consistent morphology.

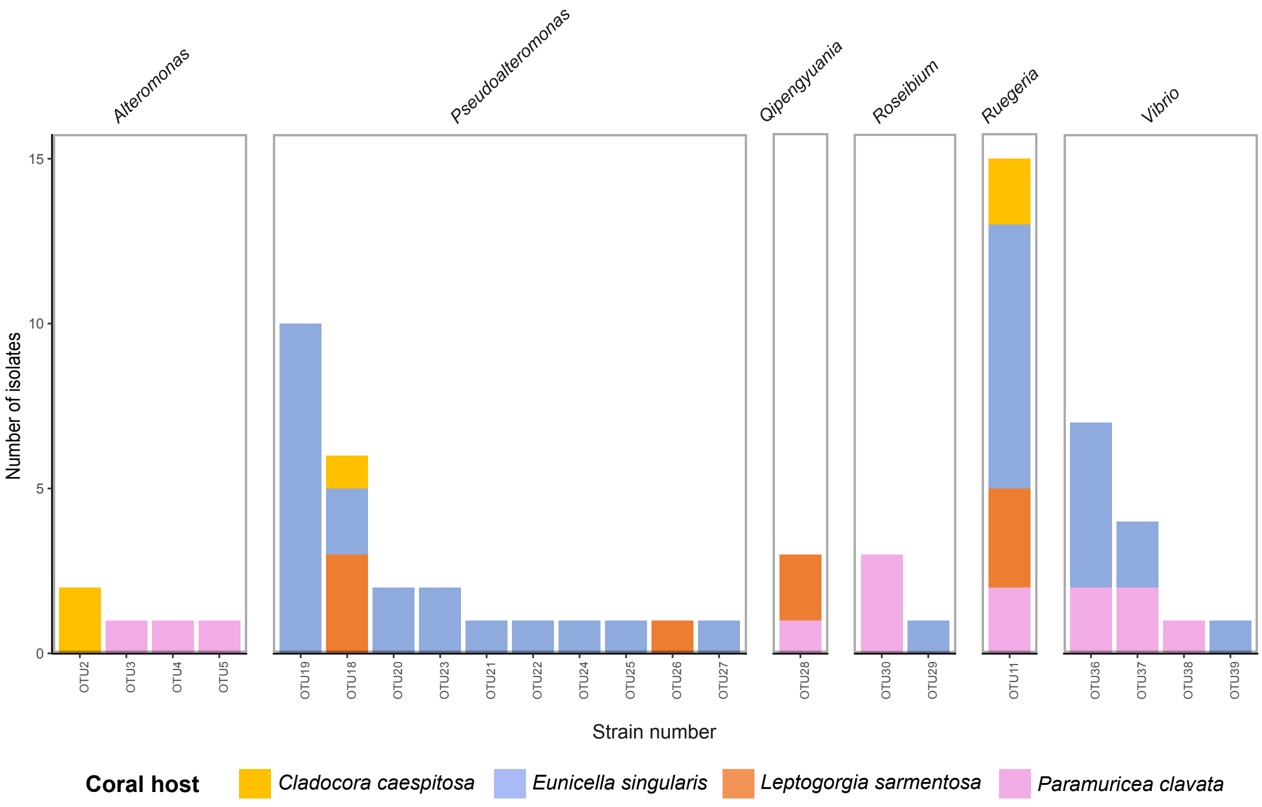


**Supplementary Figure 2. Shared and host-specific bacterial OTUs among coral hosts.** Total counts of OTUs from bacterial genera present in more than one coral species are represented in the barplot. The only strain shared by all host species belongs to the genus *Ruegeria*. In contrast, for genera such as *Alteromonas* or *Roseibium*, each coral harbors distinct bacterial OTUs.

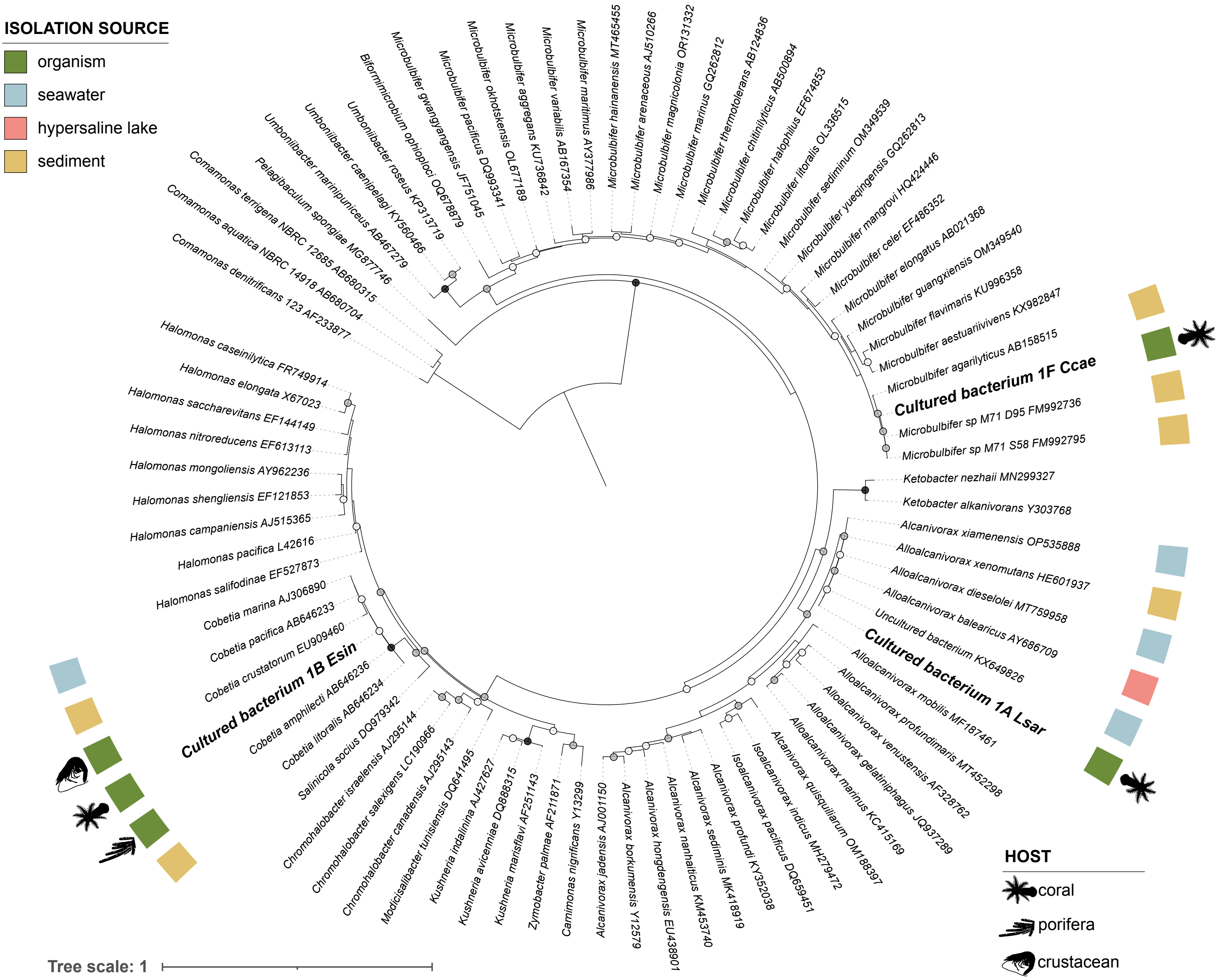


**Supplementary Figure 3. Maximum-likelihood phylogeny of Oceanospirillales and Cellvibrionales based on 16S rRNA gene sequences.** The tree includes isolates obtained in this study alongside closely related reference sequences; *Comamonas* were used as outgroup. Bootstrap support (1000 replicates) is indicated by circles at nodes (black, 100%; grey, 90–99%; white, 70–89%). Colored squares at the tips indicate isolation source, and animal silhouettes inform about the host of origin for those strains detected in organisms.

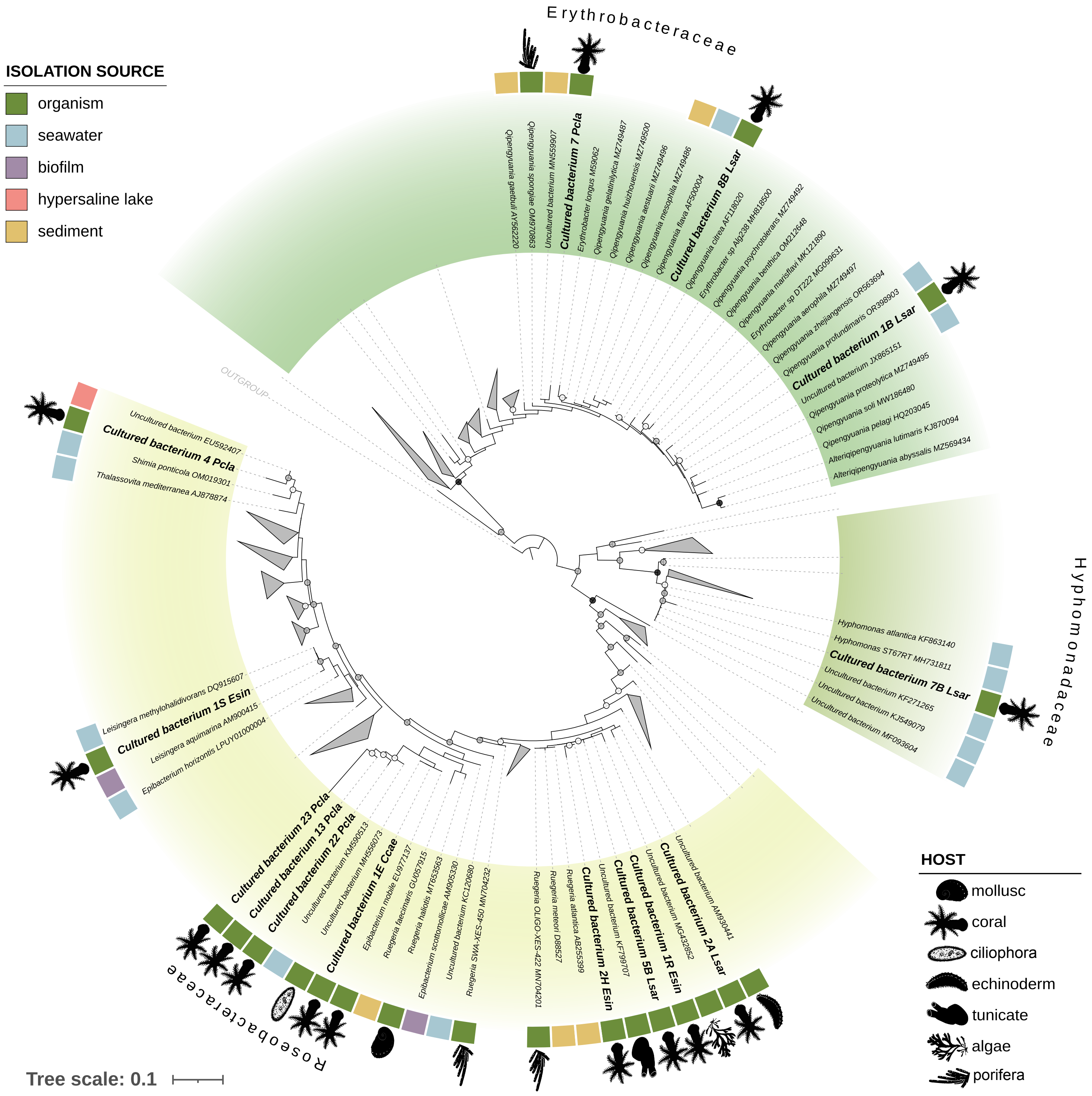


**Supplementary Figure 4. Maximum-likelihood phylogeny of Alphaproteobacteria based on 16S rRNA gene sequences.** The tree includes isolates obtained in this study alongside closely related reference sequences; representants from the classes Rhodobacterales, Hyphomicrobiales and Rhodospirillales were used as outgroup. Bootstrap support (1000 replicates) is indicated by circles at nodes (black, 100%; grey, 90–99%; white, 70–89%). Major clades are color-coded by family (Erythrobacteraceae, Hyphomonadaceae, and Roseobacteraceae). Colored squares at the tips indicate isolation source, and animal silhouettes inform about the host of origin for those strains detected in organisms.

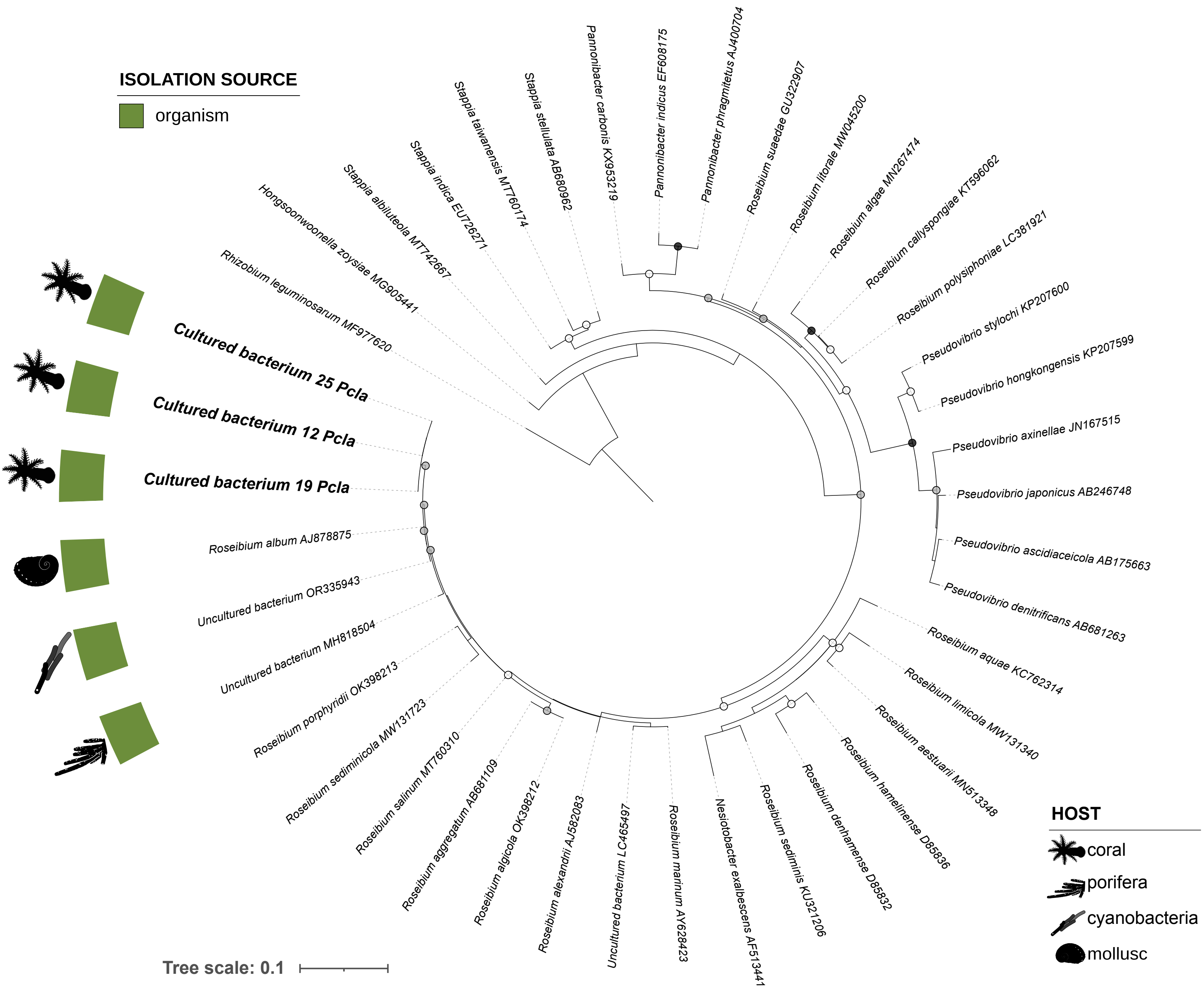


**Supplementary Figure 5. Maximum-likelihood phylogeny of Stappiaceae based on 16S rRNA gene sequences.** The tree includes isolates obtained in this study alongside closely related reference sequences; *Rhizobium leguminosarum* was used as outgroup. Bootstrap support (1000 replicates) is indicated by circles at nodes (black, 100%; grey, 90–99%; white, 70–89%). Colored squares at the tips indicate isolation source, and animal silhouettes inform about the host of origin for those strains detected in organisms.

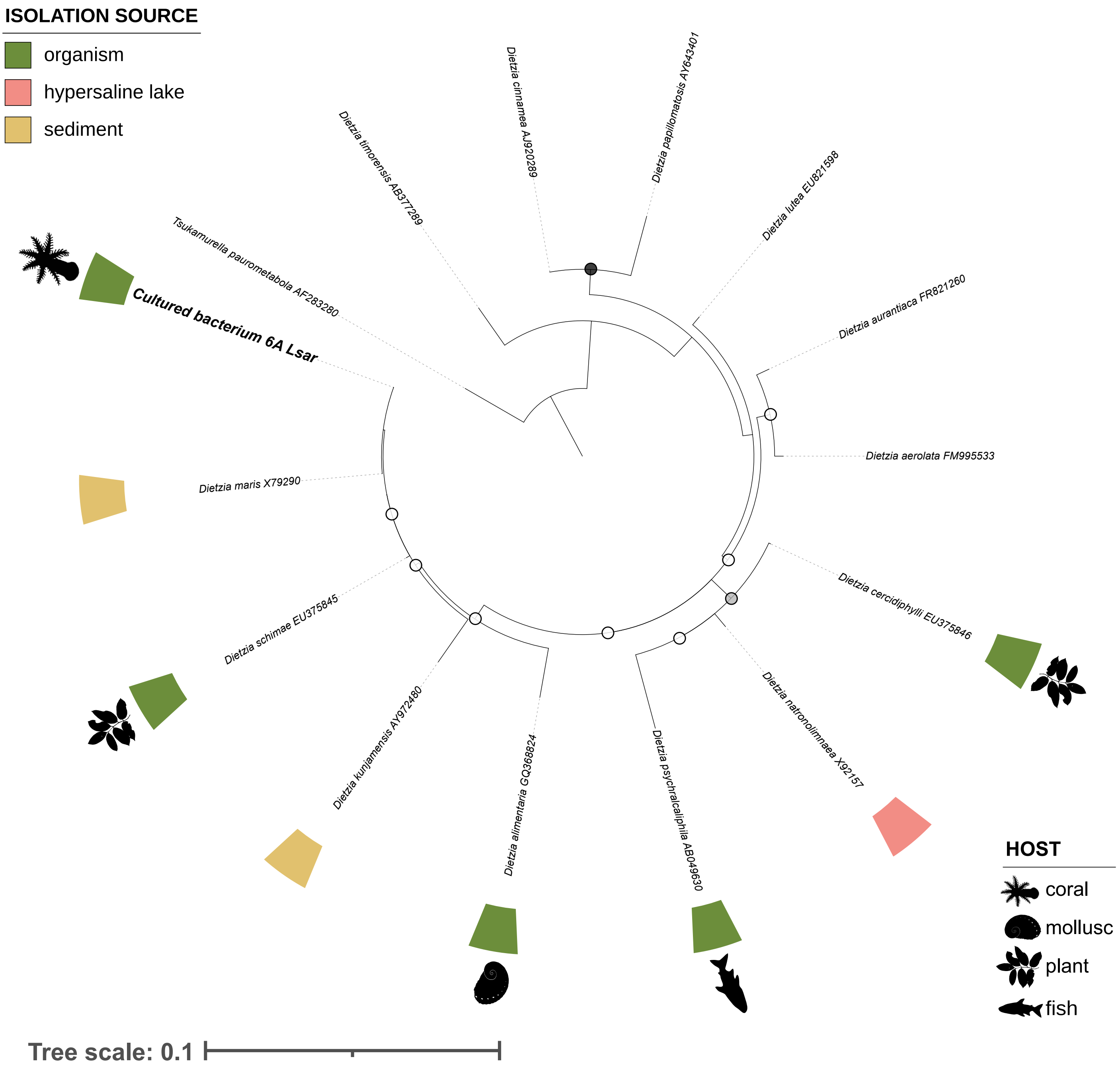


**Supplementary Figure 6. Maximum-likelihood phylogeny of Dietziaceae based on 16S rRNA gene sequences.** The tree includes isolates obtained in this study alongside closely related reference sequences; *Tsukamurella paurometabola* was used as outgroup. Bootstrap support (1000 replicates) is indicated by circles at nodes (black, 100%; grey, 90–99%; white, 70–89%). Colored squares at the tips indicate isolation source, and animal silhouettes inform about the host of origin for those strains detected in organisms.

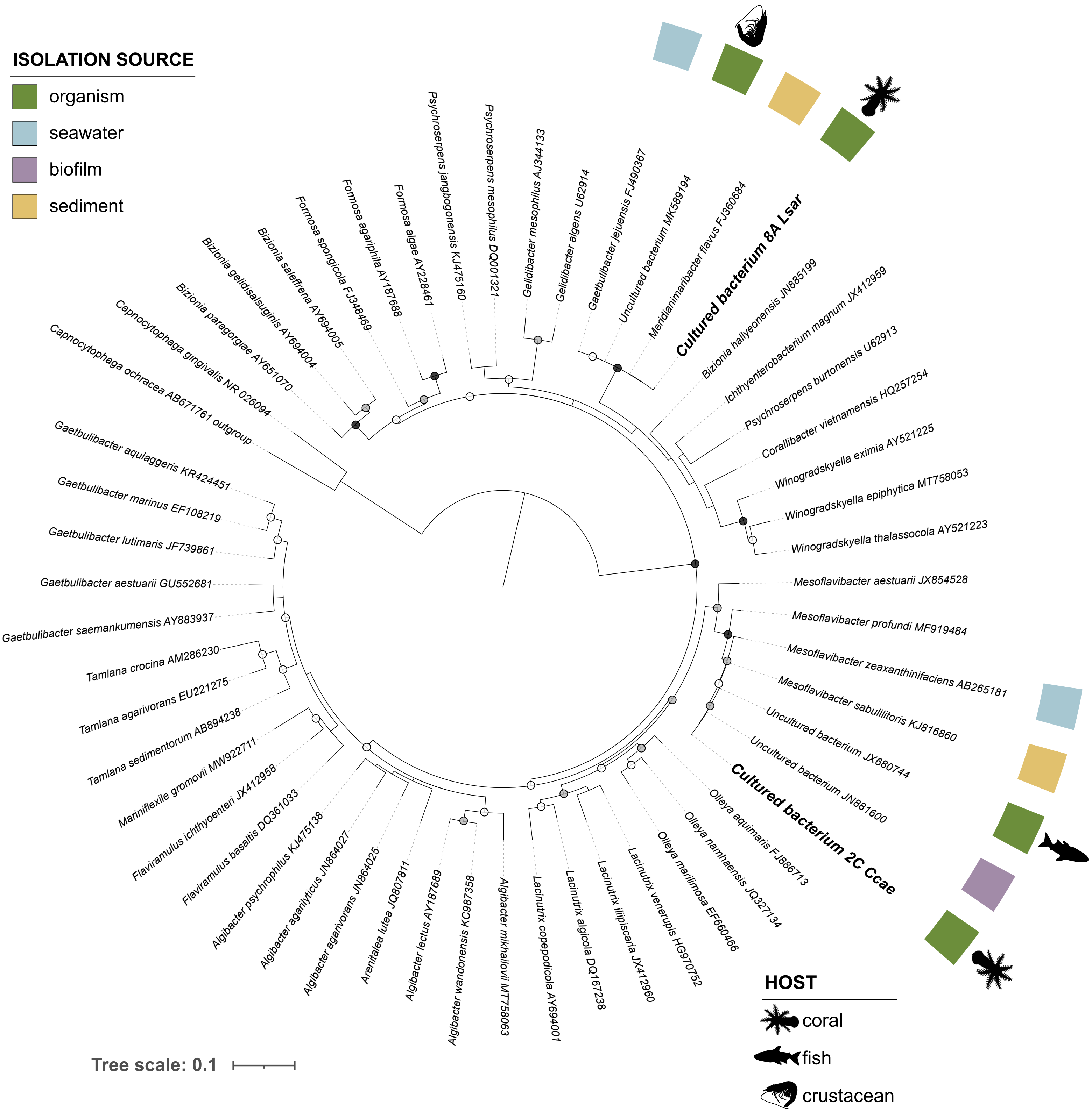


**Supplementary Figure 7. Maximum-likelihood phylogeny of Flavobacteriaceae based on 16S rRNA gene sequences.** The tree includes isolates obtained in this study alongside closely related reference sequences; *Capnocytophaga* was used as outgroup. Bootstrap support (1000 replicates) is indicated by circles at nodes (black, 100%; grey, 90–99%; white, 70–89%). Colored squares at the tips indicate isolation source, and animal silhouettes inform about the host of origin for those strains detected in organisms.
